# A basophil-specific GPCR mediates the immune response to helminth infection

**DOI:** 10.64898/2026.04.06.716327

**Authors:** Aleksander Geske, Rachel Pan, Celeste Flores, Taylor Follansbee, Nathachit Limjunyawong, Dona Ridma Harindri Sirimanna, Xintong Dong, Xinzhong Dong

## Abstract

Basophils are rare innate immune cells that contribute to type 2 immunity in allergy and parasitic helminth infection, yet the receptors governing their activation remain poorly defined. Here, we identify the Mas-related G protein-coupled receptor (GPCR) Mrgpra6 as a basophil-specific gene that mediates the host response to helminth infection. Mrgpra6 expression is restricted to basophils in blood and lung and is uniformly expressed across the population. Knockout of Mrgpra6 impairs early innate immunity to invading helminths, resulting in increased larval burden, elevated mortality, and disrupted parasite progression within the host. Transcriptional profiling of FACS-sorted basophils during helminth infection reveals how Mrgpra6 shapes the basophil transcriptional landscape needed for effective host defense. Together, these findings identify Mrgpra6 as a functional regulator of basophil-mediated anti-helminth immunity.

## Introduction

During the summer of 1878 in the city of Leipzig, Dr. Paul Ehrlich made a seminal contribution to the medical field for his discovery of mast cells and basophils, which contain dense granules positively stained by alkaline dyes. He observed that these cells were diverse in morphology and localization, ranging in size and populating connective tissue, blood, and bone marrow. Although the granules he described were not the sources of nutrition as he initially hypothesized, they instead mediate rapid and robust immune responses to allergens and parasitic helminths, forming a foundation for type 2 immunity^1,2^.

Basophils and mast cells are commonly associated together due to their shared identity as granulocytes involved in type 2 immunity. Despite these similarities, their developmental trajectories differ. Mast cells complete maturation within tissues, whereas basophils mature in the bone marrow and circulate in blood^3^. Basophils are not spatially limited and are more readily accessible for diagnostic purposes. Yet, they represent the rarest blood cell population, making up less than 1% of circulating blood leukocytes.

Large parasites such as helminths have had a profound influence on the evolution of the human immune system^4^. Today, they infect almost a quarter of the global population^5,6^. Many helminths are entirely dependent on their host for development and survival. The host immune system, on the other hand, must simultaneously promote parasite clearance and minimize tissue injury without incurring significant inflammation and further damage to vital organs like the lung^7^. It is this balance which highlights the complexity of helminth immunity and presents a challenge for clinical treatment.

Basophils are well-situated to play a role in lung anti-helminth immunity. They are important for inducing airway inflammation in models of type 2 immunity and their recruitment to the lung is often a hallmark of helminth infection^8,9^. In this early response to helminth infection, basophils are a primary source of IL4, a cytokine critical in driving type 2 immunity^10,11^. *Nippostrongylus brasiliensis* is a rodent hookworm used to model helminth infections and type 2 inflammation in mice. Mice with IL-4/IL-13–deficient T cells retain efficient *N. brasiliensis* clearance. However, concurrent basophil depletion impairs parasite elimination, highlighting the critical contribution of basophil-derived cytokines^12^. To date, the receptors mediating the basophil response to helminths are unclear.

While current understanding of mast cell biology is extensive, literature on basophil biology remains sparse in comparison. Both basophils and mast cells express the high-affinity IgE receptor (Fc□RI) and can become activated through IgE crosslinking^13^. Mast cells can also activate through a non-IgE mediated pathway, where ligand binding to the receptor Mrgprb2 results in selective degranulation of the mast cell^14,15^. In contrast, the mechanisms governing selective basophil activation and mobilization remain poorly defined.

Notably, we have previously shown that the Mrgpr family consists of multiple receptors, such as Mrgprb2, Mrgpra2a/b, and Mrgpra3, which have well-defined roles in itch and innate immunity^15–18^. However, several members remain functionally uncharacterized, raising the possibility that distinct Mrgprs may regulate basophil activation, analogous to Mrgprb2 in mast cells.

Here, we show that murine basophils specifically express the orphan GPCR Mrgpra6. CRISPR-mediated knockout of Mrgpra6 results in a significant impairment of helminth infection and a concurrent attenuation of the lung type 2 immune response. Furthermore, transcriptomic profiling of lung basophils reveals differential expression associated with basophil activation during helminth infection and identifies Mrgpra6 as a key contributor to this activation signature.

## Results

### Mrgpra6 is expressed specifically in basophils

Mrgpra6 is an orphan GPCR with uncharacterized function. We analyzed the ‘Ultra-low input’ (ULI) RNA sequencing from the Immunological Genome Project dataset, which profiles 127 highly purified populations spanning stem cells, B cells, αβ T cells, γδ T cells, innate lymphocytes, dendritic cells, macrophages, monocytes, granulocytes, mast cells, basophils, eosinophils, and stromal cells. Mrgpra6 expression was restricted to basophils, with negligible expression across all other profiled populations, including major immune lineages (Figure 1a, Data S1). Bulk RNA sequencing of FACS sorted basophils validated the robust expression of Mrgpra6 alongside basophil marker genes Mcpt8, Cd200r3, and Il4 (Figure 1b).

**Fig. 1.**
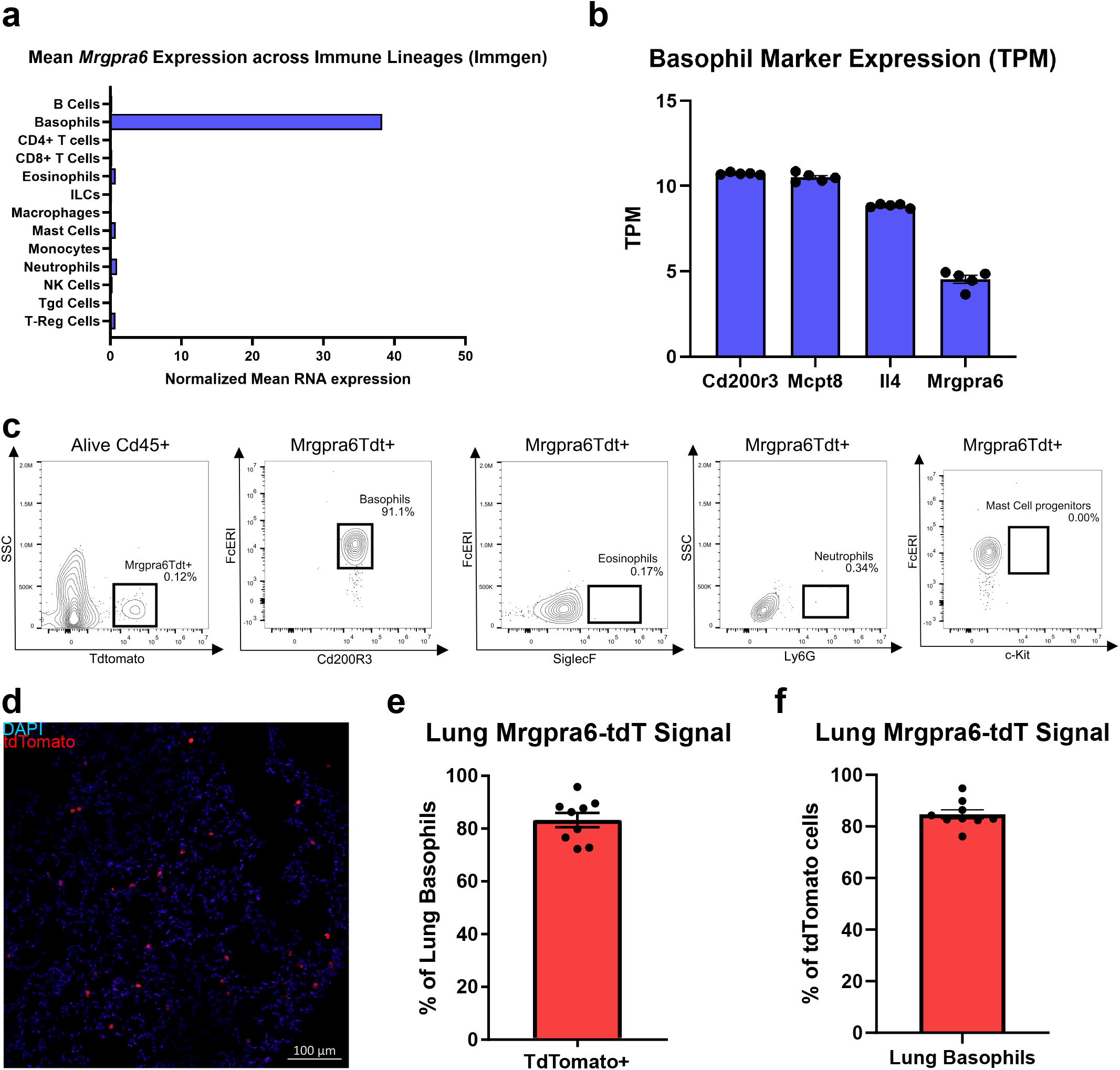
Basophils uniquely express the orphan GPCR Mrgpra6. (a) Immgen ULI RNA sequencing data showing expression of Mrgpra6 in a subset of immune cells. (b) Bulk RNA sequencing of sorted lung basophils showing normalized transcripts per million (TPM) expression of highly expressed basophil genes Cd200R3, Mcpt8, and Il4 compared to Mrgpra6. n = 5 (c) Flow cytometry gating of live, CD45□ blood cells identifying the relative contribution of basophils, eosinophils, neutrophils, and mast cell progenitors to the tdTomato□ population. (d) Representative confocal images of sections of the lung from Mrgpra6CreER; Rosa26LSL-tdTomato mice showing DAPI (blue) and tdTomato (red) expression in Mrgpra6-expressing cells. (e-f) Flow cytometric analysis of lung basophils demonstrates that ∼80% are tdTomato+, consistent with Mrgpra6 expression in this population (e). Converesly, ∼80% of lung tdTomato+ cells are Cd200r3+ FcerI+ basophils, supporting the specificity of Mrgpra6 to basophils in the lung (f). n = 9

To label Mrgpra6 expressing cells, we generated a BAC transgenic Mrgpra6^CreER^ line and crossed it with the Rosa26^LSL-tdTomato^ reporter line (Ai14), enabling tamoxifen inducible tdTomato expression in Mrgpra6-expressing cells. Given sequencing analysis showing Mrgpra6 expression uniquely in basophils, we expected tdTomato expression in the Mrgpra6^CreER^-Rosa26^LSL-tdTomato^ reporter line would be limited to basophils. Using flow cytometry, we verified that blood tdTomato expression strongly colocalized with Cd200r3+FcεrI+ basophils (92.7%). Consistent with our sequencing data, we observed only trace signal colocalizing in neutrophils (0.34%), eosinophils (0.17%), and mast cell progenitors (0.00%) (Figure 1c). Incomplete tdTomato labeling may reflect variability in CreER-mediated recombination efficiency following tamoxifen induction, although residual labeling in other cell types may reflect Mrgpra6 expression in a small subpopulation. The specificity of tdTomato labeling for basophils, consistent with the sequencing data, indicates that Mrgpra6 expression is highly restricted to the basophil population.

Basophil function is dependent on recruitment into peripheral tissues, where they interact with the surrounding microenvironment and neighboring cells. Recently, basophils have been implicated in both allergic inflammation and helminth infection within the lung^8,10^. To investigate Mrgpra6 expression in this context, we examined basophils in the lung. Lung sections depict DAPI^+^tdTomato^+^ cells throughout the alveolar walls (Figure 1d). Flow cytometry of mouse lung demonstrates that ∼80% of Cd200r3^+^ FcεrI^+^ lung basophils are tdTomato+, consistent with the expression of Mrgpra6 within this population (Figure 1e). Conversely, ∼80% of lung tdTomato+ cells are Cd200r3+ FcerI+ basophils, supporting the specificity of Mrgpra6 expression to basophils in the lung (Figure 1f). Given that other Mrgpr family members are classically expressed in peripheral sensory neurons, we examined dorsal root ganglia as an additional specificity control and observed no DAPI+tdTomato+ labeling (Supplementary Figure S1a), indicating that Mrgpra6 expression is not shared with sensory neurons. This data further validates basophil-specific Mrgpra6 expression and shows that Mrgpra6+ basophils are present in the lung, a critical site of early anti-helminth immunity.

### Mrgpra6 is required for effective anti-helminth immunity

Basophils are key contributors to type 2 immunity, with rapid IL-4 production supporting early innate immune responses and the initiation of adaptive type 2 immunity^1,10^. To investigate the role of Mrgpra6-mediated basophil function, we generated a Mrgpra6-knockout (A6KO) mouse strain using CRISPR–Cas9 and assessed female wild-type (WT) and A6KO mice in the *N. brasiliensis* model of type 2 immunity.

After subcutaneous infection with 500 infective third-stage (L3) larvae, the worms migrate via the circulation to the lung between days 0-3 post-infection, where larval passage induces acute tissue injury, inflammation, and further larval development (L4). Larvae subsequently migrate out of the lung and are swallowed, establishing their final larval stage (L5) in the small intestine. Here they mature, mate, and produce eggs between days 5–12 post-infection (Figure 2a). We first measured the extent of infection by quantifying fecal egg counts, an established endpoint of helminth infection. On days 6 and 7 post-infection, egg counts were significantly reduced in A6KO mice when compared to their WT counterparts (Figure 2b; Supplemental Figure 1b-c). To further assess disease severity during helminth infection, we conducted survival studies in mice infected with 1,000 infective L3 larvae. Survival was significantly reduced in A6KO mice (Figure 2c), with most mortality occurring during the early lung post-infection phase. These data indicate a disruption in host control of helminth infection, beginning with an altered early response in the lung.

**Fig. 2.**
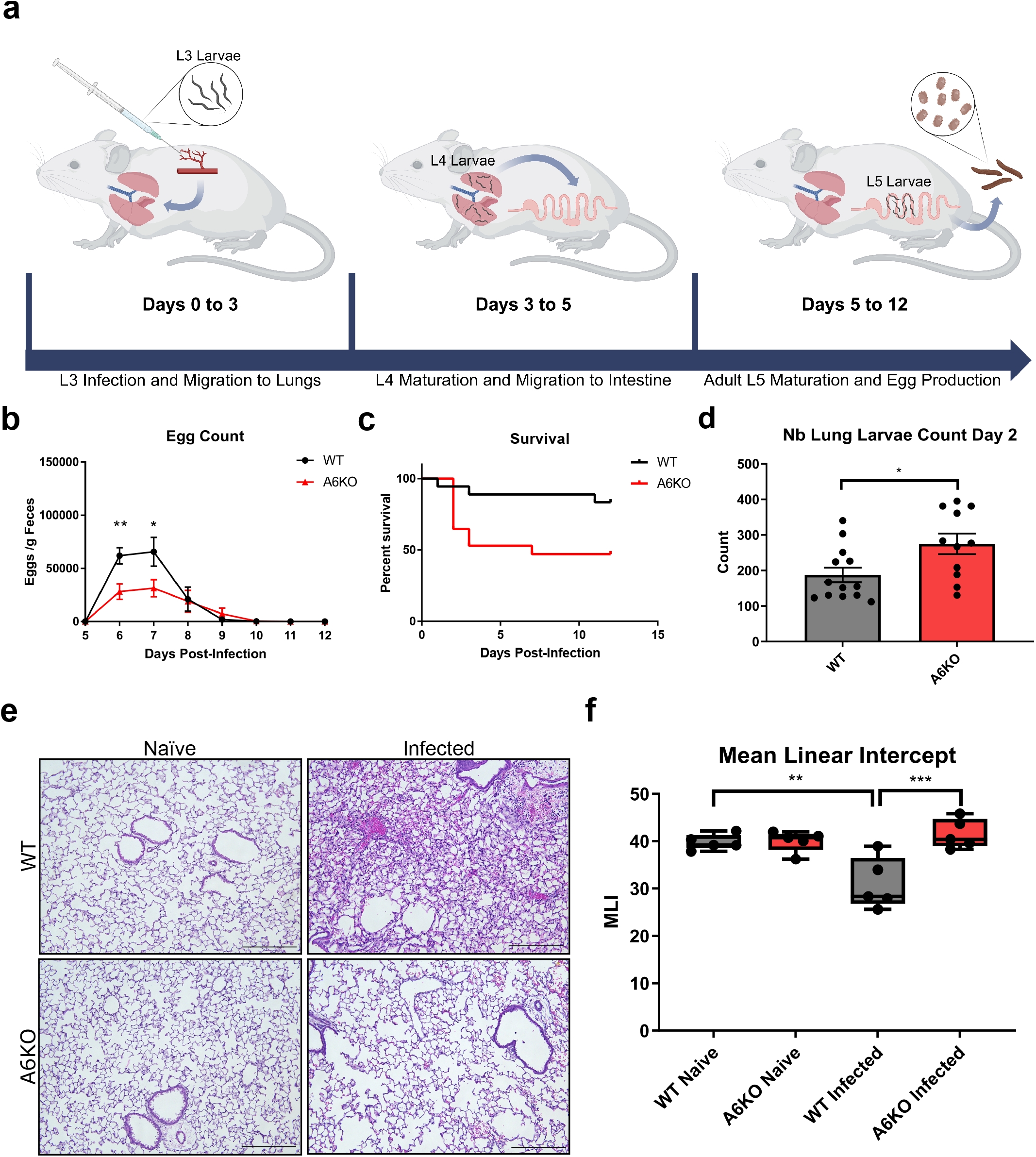
Mrgpra6 is required for effective anti-helminth immunity. (a) Schematic illustration of N. brasiliensis infection timeline. (b) Egg counts (eggs per gram of feces) from WT and A6KO mice during N. brasiliensis infection. Peak egg production occurs on days 6–7 post-infection, with significant reduced egg production in A6KO mice compared to WT. n= 14-16 (c) Kaplan–Meier survival curves for WT and A6KO mice following infection with 1,000 larvae. Survival differs significantly between WT and A6KO mice (p = 0.0246). n= 16-17 (d) Larval burden in WT and A6KO mice at day 2 post-infection, showing a significant increase in A6KO mice compared to WT. n = 11-13 (e-f) Representative H&E-stained lung sections from WT and A6KO mice under naïve and infected conditions (e), with corresponding mean linear intercept (MLI) quantification (f). WT infected mice exhibit a significantly reduced MLI compared to A6KO infected mice. Scale bar, 250µm. n = 5 Results are presented as mean ± SEM from at least three independent experiments. ∗∗p < 0.01, ∗∗∗p < 0.001, ∗∗∗∗p < 0.0001, n.s., not significant by two-way ANOVA with Šídák’s multiple comparisons test (b and e), log-rank test (c), two-tailed unpaired Student’s t test (d).

Larval migration through the lung can cause substantial tissue damage, resulting in edema, inflammatory cell recruitment, hemorrhage, and, in severe cases, respiratory failure. To assess parasite burden in the lung, whole lungs were harvested and viable larvae were enumerated at day 2 post-infection, the peak of lung larval burden. We observed significantly higher larval burden in A6KO compared to WT controls, consistent with the survival outcomes (Figure 2d). The combination of increased lung larval burden and reduced intestinal egg counts suggests an impairment in the helminth life cycle progression within the host. Increased larval presence in the lung suggests defective larval migration out of the lung. Proper helminth development is closely intertwined with the host environment, underscoring the importance of examining infection-induced changes in lung tissue that may disrupt helminth infection.

To investigate the lung response to infection, we hematoxylin and eosin (H&E) stained lung sections from mice at day 3 post-infection, corresponding to the peak of innate immune responses prior to larval exit from the lung. This revealed prominent inflammatory infiltrates, edema, and hemorrhage in infected WT mice, features characteristic of the lung response to acute lung injury. In contrast, these pathological features appeared less pronounced in A6KO mice (Figure 2e; Supplemental Figure 1d-g). Excessive cellular and fluid influx results in reduced airspace in the lung. Accordingly, we quantified lung architecture using mean linear intercept (MLI), a measure of alveolar airspace and indirect measure of acute lung injury, where a lower MLI shows lower airspace volume^19^. MLI was reduced in WT but not A6KO mice after infection (Figure 2f), indicating a differential response to acute lung injury and supporting observations of reduced inflammatory infiltration and edema in A6KO mice (Figure 2e). Collectively, these findings strongly support a key role of Mrgpra6 in the early lung-phase response to *N. brasiliensis* infection, with knockout of Mrgpra6 resulting in defective larval clearance from the lung and altered tissue responses in mice.

### Type 2 Immunity is impaired in the absence of Mrgpra6

Given the differential lung responses to *N. brasiliensis* infection observed in WT and A6KO mice, we next examined the lung immune response to better define the underlying immunological differences. Using flow cytometry, we quantified total CD45□ leukocytes and CD200R3□FcεRI□ basophils in perfused lungs at day 3 post-infection, restricting analysis to extravascular cells. *N. brasiliensis* infected WT mice exhibited a substantial increase in leukocyte and basophil recruitment compared to non-infected mice, an effect which was significantly attenuated in A6KO mice (Figure 3a).

**Fig. 3.**
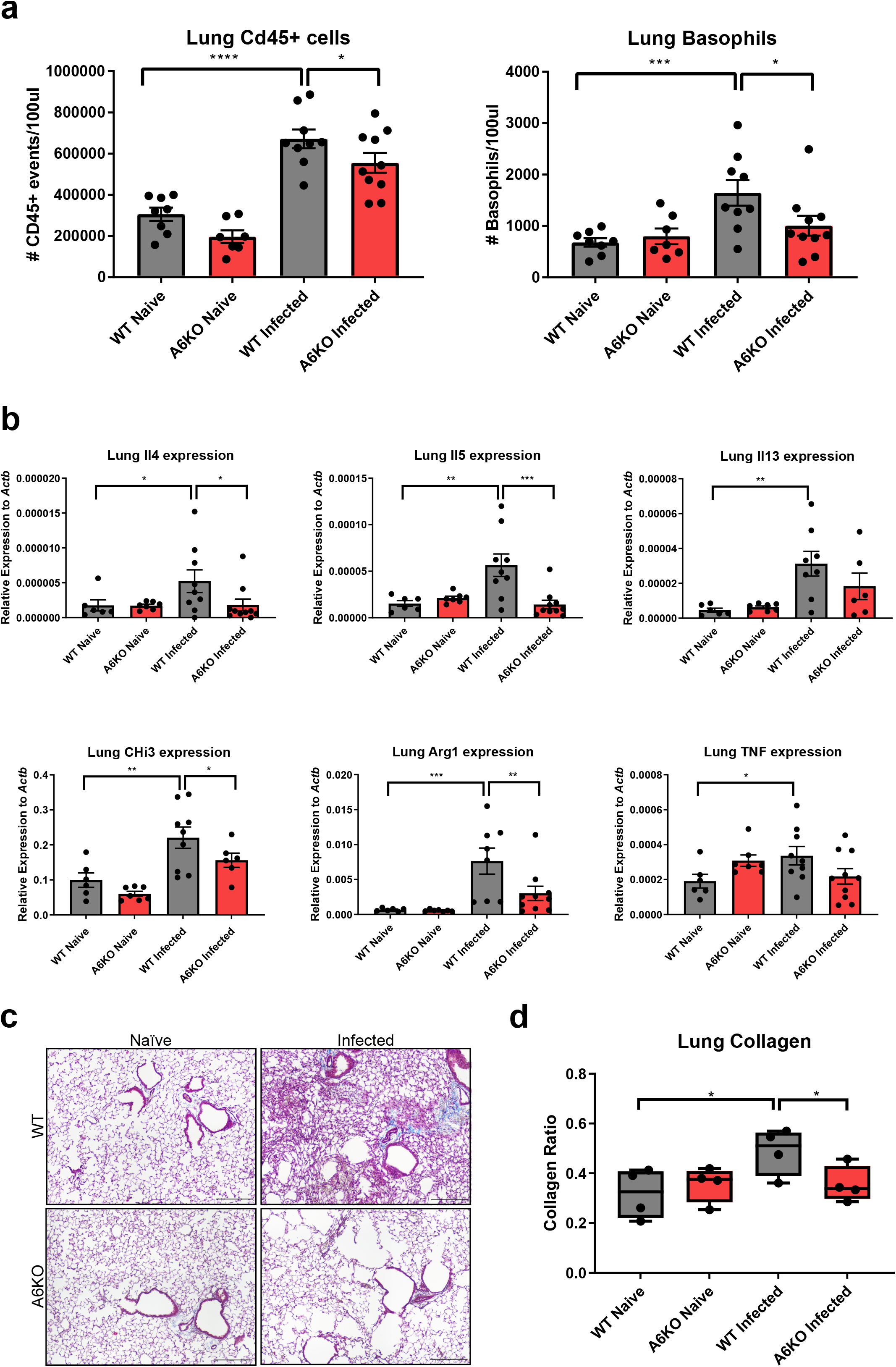
Reduced Type 2 Immunity in A6KO female mice. (a) Flow cytometric analysis of total CD45□ leukocytes and basophils in lungs at day 3 post–N. brasiliensis infection. Infection induces robust CD45□ leukocyte recruitment in WT mice compared to A6KO mice. Basophil recruitment is significantly increased in infected WT lungs but not A6KO mice. n = 7-10 (b) qPCR quantification of lung mRNA expression for genes associated with Th2 immunity. Expression of Il4, Il5, Il13, Chil3, and Arg1 is significantly increased following N. brasiliensis infection in WT mice, whereas expression is reduced or absent in A6KO mice. Tnf expression shows a modest infection-associated increase that in WT mice. n = 6-10 (c-d) Representative Masson’s trichrome–stained lung sections from WT and A6KO mice under naïve and infected conditions (c), with corresponding collagen quantification (d). WT mice display a significant infection-associated increase in lung collagen compared to A6KO mice. Images were acquired and processed identically for brightness and contrast. Scale bar, 250µm. n = 4 Results are presented as mean ± SEM from at least three independent experiments. ∗∗p < 0.01, ∗∗∗p < 0.001, ∗∗∗∗p < 0.0001, n.s., not significant by one-way ANOVA.

Next, we assessed the expression of key type 2 markers and cytokines in the lung using qPCR. We found that WT, but not A6KO, mice exhibited significant upregulation of type 2 cytokines *Il4, Il5*, and *Il13* at day 3 post-infection (Figure 3b). Alveolar macrophages play an important effector function in innate helminth immunity in the lung. Alternative M2 activation by type 2 cytokines induces expression of *chitinase-like 3* (*Chil3*) and *arginase 1 (Arg1*)^20^. We observed a significant increase in Chil3 and Arg1 expression following infection in WT mice, whereas A6KO mice showed no such increase (Figure 3b). Together, these findings indicate an impaired early immune response in A6KO mice, consistent with the attenuated response to acute lung injury observed during infection.

Collagen deposition driven by myofibroblast recruitment and activation is initiated during the early innate immune response and, if sustained, can result in pathogenic fibrosis when collagen synthesis exceeds collagen degradation. However, early collagen synthesis is a critical and protective component of tissue repair and wound healing^21,22^. Basophils have been shown to trigger myofibroblast activation in an *Il4* dependent manner^23^. For this reason, we used Masson’s Trichrome staining to quantify collagen deposition in the early immune response to *N. brasiliensis*. Representative Masson’s trichrome staining at day 3 post-infection reveals prominent collagen staining (blue) in WT infected mice, reflecting collagen expansion and increased collagen density along alveolar walls and surrounding blood vessels (Figure 3c; Supplemental Figure 2). Quantification of collagen staining normalized to tissue area demonstrates a significant increase in collagen signal in WT mice, but not in A6KO mice (Figure 3d). This data identifies Mrgpra6 as a key regulator of early collagen deposition in the lung following helminth infection.

### Basophil transcription is dependent on Mrgpra6 in response to *N. brasiliensis* infection

Due to the low abundance of basophils relative to other immune cell populations, the transcriptional changes of basophils during helminth infection are poorly defined. To directly assess how basophils respond to *N. brasiliensis* infection and how this response is altered by knockout of Mrgpra6, we performed bulk RNA sequencing on FACS-sorted lung basophils from individual WT and A6KO mice under naïve and day 3 post-infection conditions (Figure 4a).

**Fig. 4.**
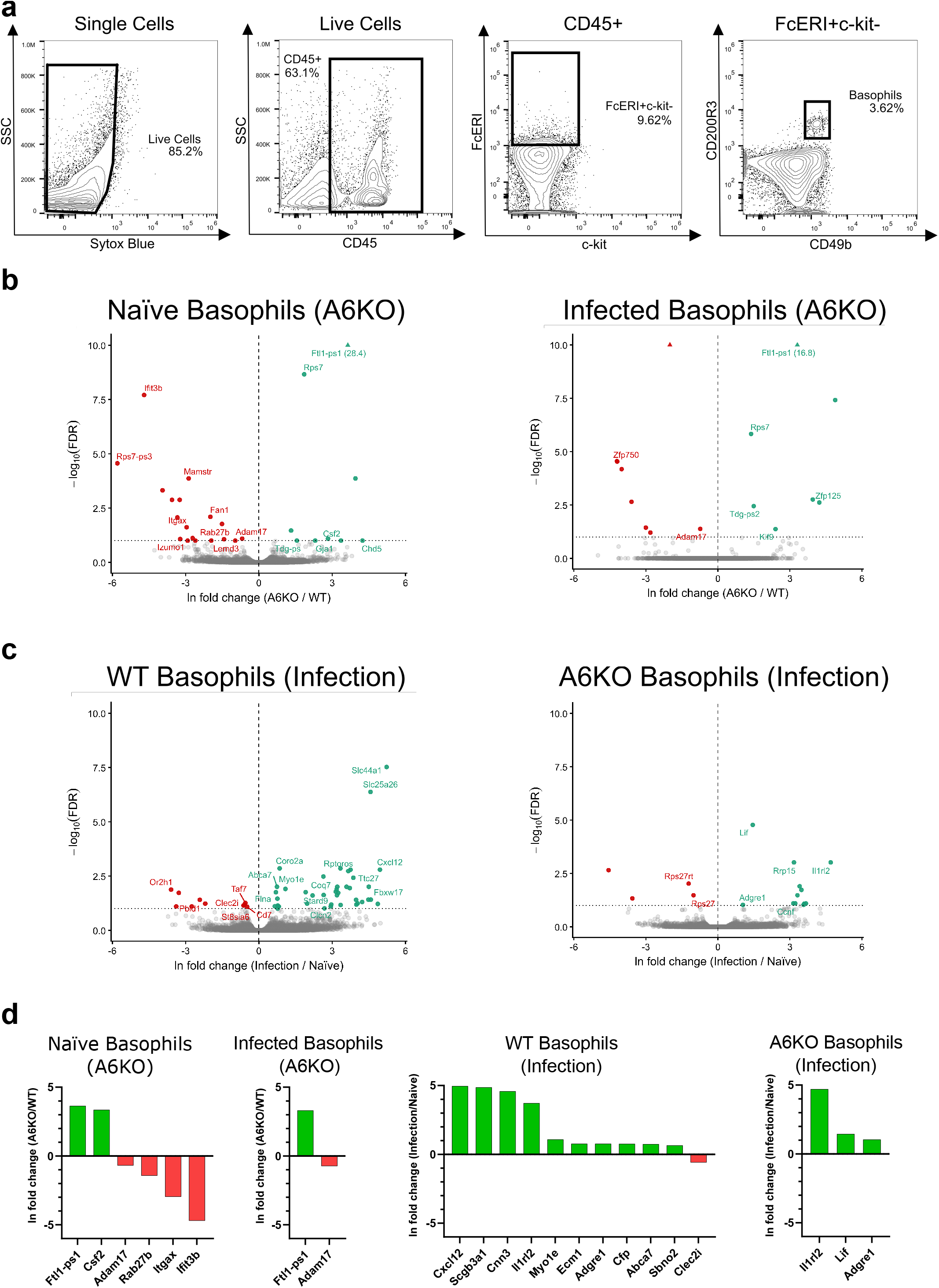
A6KO mutes basophil response to N. brasiliensis infection. (a) Flow cytometry gating strategy for basophil sorting. Single cells were gated followed by live cells (Sytox Blue-). CD45+ leukocytes were then selected and basophils identified as FcεRI+ c-kit- CD200R3+ CD49b+ cells. Cells within the basophil gate were used for FACS sorting. (b) Volcano plots showing significantly up-regulated (green) and down-regulated (red) genes in naïve and N. brasiliensis infected basophils. Plots illustrate expressional changes due to Mrgpra6 knockout (A6KO). Points whose −log□□(FDR) exceed the axis range are denoted with a triangle and the given value in parentheses next to the gene label. (c) Volcano plots showing significantly up-regulated (green) and down-regulated (red) genes in WT and A6KO basophils. Plots illustrate expressional changes due to N. brasiliensis infection (Infection). (d) Bar graphs showing effect sizes (β) (ln fold change) for selected genes in basophils due to Mrgpra6 knockout (A6KO) or N. brasiliensis infection (Infection). Differentially expressed genes were defined using an FDR-adjusted q-value < 0.10, n = 5 individual mice per genotype per condition (WT Naïve, A6KO Naïve, WT Infected, A6KO Infected)

We performed differential expression analysis to define genes regulated by *N. brasiliensis* infection in basophils and to determine how these responses are altered by Mrgpra6 knockout (Figure 4b-c; Supplemental Figure 3, Data S2). Basophil identity was further supported by robust expression of the hallmark basophil cytokines *Il4, Il6*, and *Il13* across all groups. However, expression of these genes did not significantly differ between genotypes or infection conditions, suggesting Mrgpra6 does not strongly regulate expression of these cytokines in this model (Data S2). Under naïve conditions, Mrgpra6 knockout resulted in 24 significantly differentially expressed genes in basophils (Figure 4b, Supplemental Figure 4a). Among these, A6KO basophils exhibited significant downregulation of *Adam17, Rab27b, Itgax (Cd11c)*, and *Ifit3b*, genes involved in mediator release and cytokine signaling (*Adam17, Rab27b*), cell adhesion and migration (*Itgax*), and innate immune activation (*Ifit3b*) (Figure 4d; Supplemental Figure 4). Notably, *Csf2*, which encodes the pleiotropic cytokine GM-CSF, is upregulated in A6KO naïve basophils.

Following *N. brasiliensis* infection, WT basophils exhibited substantially broader transcriptional changes, with 51 infection-associated DEGs compared to 16 DEGs in A6KO basophils (Figure 4c). Notably, these infection-associated transcriptional changes are largely upregulation, with 41 DEGs in WT basophils and 13 DEGs in A6KO basophils. The reduced number, and magnitude, of infection-induced DEGs in A6KO basophils indicates a reduced transcriptional response to infection. Following *N. brasiliensis* infection, WT basophils have a significant upregulation of immune-associated genes compared to A6KO basophils. Notably, *Il1rl2* and *Lif*, which encode an Il36 cytokine receptor and a Il6-family cytokine, respectively, remain upregulated in A6KO basophils. *Il1rl2* is similarly upregulated in WT basophils, whereas *Lif* shows a weaker, non-significant upregulation. This reduced upregulation of immune-associated genes in A6KO basophils is consistent with earlier observations and provides functional context for the global transcriptional changes observed in A6KO basophils.

Collectively, these findings indicate that Mrgpra6 regulates the basal transcriptional state of basophils, with knockout of Mrgpra6 attenuating the basophil transcriptional response to helminth infection.

## Discussion

Here, we identify Mrgpra6 as a basophil-specific marker in mice and demonstrate that basophil-intrinsic Mrgpra6 is required for effective anti-helminth immunity. Using A6KO mice, we uncover defects in both host immune responses and helminth progression during infection. To our knowledge, we provide the first transcriptional analysis of basophils during helminth infection, defining the gene expression profile that governs their response and revealing how Mrgpra6 shapes this transcriptional state.

Currently established basophil markers lack clear functional characterization. Mcpt8, a serine protease specifically found in basophil granules, can elicit inflammation upon intradermal injection. However, how and when basophils activate and release Mcpt8 is in need of further study^24^. Human and mouse basophils highly express CD200R upon activation, though the physiological significance of this upregulation is not well defined^25^. Our findings identify Mrgpra6 as a basophil-specific GPCR with direct functional relevance in anti-helminth immunity.

Helminth infections are designated by the World Health Organization as neglected tropical diseases (NTDs), highlighting a mismatch between their widespread prevalence and the scientific attention they receive ^26^. The biology underpinning host–helminth tolerance and clearance remains incompletely understood. Here, we highlight the lung as an important checkpoint in helminth immunity.

The lungs are a tightly regulated immune environment, where inflammation must be limited to prioritize the health and fitness of the host. This regulatory balance may be particularly important for helminths whose life cycle requires migration through the lung; however, the mechanisms underlying this dependency remain unclear^27^. Helminth development may depend on biochemical cues within tissue^27^. Indeed, multiple lines of evidence suggest that helminths can exploit host immune cues for their development. IL-5 enhanced *Litomosoides sigmodontis* development^28^, IL-4/STAT6 signaling in eosinophils was shown to be required for normal *Trichinella spiralis* larval growth^29^, and RELMα-expressing macrophages maintained *N. brasiliensis* larval motility and viability relative to RELMα-deficient macrophages^30^. *N. brasiliensis* necessarily migrate through the lung to progress from larval stages L3 to L4, suggesting a similar environmental dependency on the tissue environment and subsequent type 2 response. However, *N. brasiliensis* developmental biology remains incompletely characterized, and the host biochemical cues that govern this process are undefined. Here, we illustrate this delicate balance and show that an attenuated type 2 immune environment is associated with impaired helminth progression at multiple stages of the life cycle. Mrgpra6 knockout mice show increased larval burden in the lung, consistent with delayed or impaired larval migration. Moreover, Mrgpra6 knockout mice exhibit reduced egg counts, a critical impairment of helminth reproductive output and disruption of the helminth life cycle.

Basophils are known to be recruited during helminth infection, though their function in anti-helminth immunity remains less clear. Prior work emphasizes their production of IL-4 and contribution to the initiation of adaptive type 2 immunity^10^. We define a niche for this rare granulocyte in the innate response to helminths. Knockout of Mrgpra6 reduces type 2 immunity in response to *N. brasiliensis* infection, with respect to overall leukocyte recruitment, basophil recruitment, and the expression of key type 2 cytokines and markers. Furthermore, knockout of Mrgpra6 results in reduced early collagen deposition, consistent with an impaired tissue repair response associated with type 2 immunity. Together, our findings highlight the role of basophils in the early innate response to helminth infection in the lung, contributing to effective type 2 immunity during the critical period of lung worm clearance and tissue repair.

With findings from lung basophil RNA sequencing, we shed light on transcriptional changes in response to helminth infection and the role Mrgpra6 plays in these changes. In naïve lung basophils, expression of *Adam17, Rab27b, Itgax*, and *Ifit3b* are significantly reduced. *Adam17*, the integral sheddase, proteolytically cleaves membrane bound immune mediators such as TNFα and *Il6R*, enabling their release into tissue^31^. *Rab27b* regulates degranulation in mast cells, potentially having similar functions in basophil degranulation^32^. *Itgax*, also known as *Cd11c*, mediates leukocyte interactions including cell adhesion and complement activation^33^. *Ifit3b* is homologous to human *IFIT3*, an interferon-stimulated gene regulated by the JAK–STAT pathway, which may be attenuated in Mrgpra6 KO basophils^34^. Together, the downregulation of this group of genes suggests a deficit in basophil function and further implicates Mrgpra6 as an important regulator of basal basophil activity. Furthermore, WT basophils upregulate a host of immune genes in response to helminth infection, whereas this response is largely absent in Mrgpra6 knockout basophils. This suggests that Mrgpra6 enables a broader transcriptional response to helminth infection. Given that basophils are established mediators of type 2 immunity, this muted transcriptional response may contribute directly to the attenuated type 2 environment, or indirectly through downstream effects on other anti-helminth effector populations, including eosinophils and macrophages. This attenuated response may in turn underlie the impaired larval clearance and helminth life cycle progression described above.

The most significant differentially expressed gene in A6KO basophils is Ftl1-ps1, potentially representing Ftl1 due to significant sequence overlap between the Ftl1-ps1 and Ftl1 transcripts. FTL, the human ortholog of murine Ftl1, was recently identified as the most significantly differentially expressed marker of a transcriptionally distinct, circulating basophil population^35^. Given increased expression of genes which have been shown to downregulate with basophil maturity ^35–37^, FTL expressing basophils may represent a functionally immature basophil state in humans. The upregulation of Ftl1-ps1 in A6KO basophils raises the hypothesis that Mrgpra6 loss skews basophils toward an immature transcriptional state, potentially impairing capacity to respond to infection. The function of FTL/Ftl1 in basophils is, however, unknown. This interpretation remains speculative and requires further investigation.

In human basophils, MRGPRX2 expression is detected^38^, potentially making it the human ortholog to Mrgpra6, alongside the established mouse ortholog Mrgprb2. While MRGPRX2 expression in unstimulated basophils is contested, basophil MRGPRX2 surface expression appears to be upregulated following basophil stimulation^39,40^. How MRGPRX2 functions in human basophils, and whether it meaningfully contributes to helminth immunity, is unclear. Further characterization of this candidate human ortholog is needed.

Overall, these findings support a model where Mrgpra6 contributes to the basal function of basophils, influencing their transcriptional state and ability to regulate early innate immune responses in the lung. Disruption of this pathway significantly attenuates the host response during the critical window of larval clearance from the lung, resulting in increased host mortality and impaired helminth life cycle progression. These findings position basophils as non-redundant mediators of innate anti-helminth immunity and reveal Mrgpra6 as a regulator of early type 2 immunity in the lung.

## Materials and Methods

### Experimental Model and Mouse Details

#### Mice

C57Bl/6J, *Mrgpra6*-CRISPR-KO, and Mrgpra6-CreER mice were bred under specific pathogen-free conditions in the Johns Hopkins University School of Medicine Animal Facility and weaned at 3 weeks of age. Mrgpra6-CreER; Rosa26-lsl-tdTomato reporter mice were obtained by crossing Mrgpra6-CreER animals with Rosa26-LoxP-STOP-LoxP (lsl)-tdTomato animals (Jackson labs). Mice were kept in community cages (4-5 mice per cage) at light period of 12 h and fed water and mouse chow *ad libitum*. 8-12 weeks old, age-matched female mice were used for experiments. All animal procedures were conducted in accordance with institutional guidelines and with approval from the Animal Care and Use Committee at the Johns Hopkins University School of Medicine.

#### Helminth Passage, Culture, and Collection

Infectious stage-three *N. brasliensis* larvae were harvested from fecal culture using a Baermann apparatus. Mice were infected subcutaneously into the dorsal neck (scruff) region with 1000 larvae for survival study and 500 larvae for all else. To make fecal cultures, *N. brasliensis* larvae were passaged through mice by inoculating them with 500 larvae per mouse. At day 6 post infection, feces from infected cecum and large intestine were collected into tap water, mixed with activated charcoal (Sigma-Aldrich) and peat moss (Miracle-Gro) (1:1), and placed on a moist, lint-free ChemWipes (Kimberly-Clark).

### Method Details

#### Generation of Mrgpra6 KO and Mrgpra6Cre-ER mice

*Mrgpra6* KO were generated by the Johns Hopkins University School of Medicine Transgenic Core Laboratory using standard CRISPR/cas9 methods. Specifically, mice were generated using CRISPR with gRNA targeting the second exon of the Mrgpra6 coding region. The KO allele carries a 299 base pair deletion starting with the sequence TAGTCGGGCTGACAGGA and ending with the sequence GGTATCGCTGCCACCGCCCA. Successful deletion was verified by PCR and Sanger sequencing. Mrgpra6CreER BAC mice were generated by the Gene Targeting & Transgenic Facility at Janelia Farm (Ashburn, VA) using BAC clone RP23-208C7 (C57BL/6J-derived library).

#### Mouse Genotyping

Primers and annealing temperatures were as follows:

Mrgpra6Cre-ER – CACCTCAACCTCGGCAAC (F), CGTTCACCGGCATCAACGT (R); annealing temperature 57°C

Mrgpra6 KO – GTGTTCATAGTGAAGGCCTC (F), GGAAGTTCATTGCTAGACACC (R); annealing temperature 57°C

tdTomato – AAGGGAGCTGCAGTGGAGTA (WT-F), CCGAAAATCTGTGGGAAGTC (WT-R), GGCATTAAAGCAGCGTATCC (MUT-F), CTGTTCCTGTACGGCATGG (MUT-R); annealing temperature 61°C.

#### Helminth egg count and viable lung larvae recovery

For egg counts, feces was collected daily in 1ml of water and vortexed intermittently for 1 hour. Each sample was then diluted in NaCL solution (35.7g/100ml) at either 2x or 5x dilutions, causing eggs to float up. Eggs were then counted in the McMaster Egg Slide Chamber (Electron Microscopy Sciences). For lung larvae counts, lung tissue was minced, placed on cheesecloth, and suspended in PBS warmed to 37°C for 6 hours. Viable worms collect to the bottom of the tube and are counted on the Nikon Ti2 inverted microscope.

#### Histology

Lungs inflated with 4% PFA were perfused with PBS and placed into 4% PFA prior to submission to The Johns Hopkins University Oncology Tissue and Imaging Service (OTIS) Core Laboratory where they were embedded in paraffin, sliced into μm sections, and stained with hematoxylin and eosin (H&E). Masson’s trichrome staining was performed using the Trichrome Stain Kit (Abcam) according to manufacturer’s instructions. Microscopy was performed using the Nikon Ti2 inverted microscope. For fluorescence labeling, lungs inflated with 4% PFA were perfused with PBS, placed into 4% PFA at 4°C overnight, then placed in 30% sucrose at 4°C overnight. Lungs were then stored in optimal cutting compound (OCT) at -80°C until sectioned with a cryostat at 20uM sections onto solids. Sections were washed with PBST (0.3% Triton X-100), then mounted with Fluoromount-G with DAPI (ThermoFisher). Microscopy was performed using the Zeiss LSM700 Confocal microscope.

#### Lung Processing and Flow Cytometry

Prior to lung harvest, mice were perfused with PBS via the right ventricle to remove intravascular blood cells. Lungs were minced and digested in 5mls RPMI containing 0.1mg/ml Liberase TL (Roche) and 30ug/ml DNase I for 30 minutes at 37C. Single cell suspensions were generated by passing digested lung through a 70um cell strainer (VWR). Live versus Dead cells were staining using Live/Dead Fixable Aqua Dead Cell Stain Kit (Thermo Fisher) for standard flow cytometry and Sytox Blue (Thermo Fisher) for FACS. Cells were treated with CD16/CD32 Fc Block (S17011E, Biolegend) for 10 min before incubation with antibodies. For flow cytometry and FACS sorting of lung, the following antibodies were used: CD45-APC/Cy7(Biolegend), CD117-c-kit-BV605(Biolegend), FceRI-PE/Cy7(Biolegend), CD200R3-APC(Biolegend), Cd49b-PerCP-Cy5.5(Biolegend). For flow cytometry of blood, the following antibodies were used: SiglecF BB515 (Biolegend), Ly6G BV421 (Biolegend), CD45-APC/Cy7(Biolegend), CD117-c-kit-BV605(Biolegend), FceRI-PE/Cy7(Biolegend), CD200R3-APC(Biolegend), Cd49b-PerCP-Cy5.5(Biolegend). Data were collected using a CytoFLEX LX (Beckman Coulter) and analyzed using FlowJo (BD). Basophils were gated as CD45^+^c-kit^-^FceRI^+^CD200R3^+^Cd49b^+^.

#### RNA extraction and quantification

Snap-frozen left lung tissue was homogenized using the soft tissue homogenizing kit (Precellys) beads RLT buffer via the Minilys Homogenizer (Berlin). RNA was extracted using the Rneasy® Plus Micro Kit (Qiagen) and mRNA was reverse-transcribed into cDNA using a High-Capacity cDNA Reverse Transcription Kit (Applied Biosystems). Quantitative PCR (qPCR) was performed using TaqMan™ Fast Advanced Master Mix (Applied Biosystems) and Taqman gene expression probes (Thermo Fisher) corresponding to each gene of interest on a StepOnePlus Real-Time PCR System (Applied Biosystems). Gene expression was normalized to *Actb* using the ΔCt method. The same samples were measured for multiple mRNAs.

#### Fluorescence-actvated cell sorting (FACS)

Basophils were isolated from single-cell lung suspensions, prepared as previously described. Lung samples were incubated with CD16/CD32 Fc Blocker (Biolegend) for 10 min in FACS buffer (PBS supplemented with 2% fetal bovine serum). Surface staining was performed for 30□min at 4°C with the antibodies listed prior. Dead cells were stained with SYTOX Blue (Thermo Fisher). Cell sorting was performed on the Sony MA900 cell sorter (Sony Biotechnology) with a 100 μm nozzle. Basophils were gated as CD45^+^c-kit^-^ FceRI^+^CD200R3^+^Cd49b^+^.

#### RNA sequencing

Sorted basophils were extracted from WT and Mrgpra6 KO naïve or infected lungs (72 hours post subcutaneous injection of 500 larvae) using the Arcturus™ PicoPure™ RNA Isolation Kit (Applied Biosystems), with each sample consisting of RNA extracted from individual mouse whole lung. 5 samples per condition were submitted to the Johns Hopkins Single Cell & Transcriptomics Core. Libaries were constructed using the SMART-seq mRNA LP library prep and sequenced on an Illumina NovaSeq X system. Analysis was performed using kallisto (33) for alignment and sleuth (34) for differential expression analysis. Given the exploratory nature of this analysis and the inherent limitations on statistical power imposed by low RNA input from a rare, sorted cell population, differentially expressed genes were defined using a relaxed FDR-adjusted q-value threshold of < 0.10 to reduce the risk of overlooking true positives at the expense of an increased false discovery rate.

#### Quantification and Statistical Analysis

Image quantifications were performed using ImageJ (NIH). All statistical analyses were performed using Prism 9 (GraphPad). Single comparisons were made using two-tailed unpaired Student’s t test. Data for multiple comparisons were analyzed using one-way or two-way ANOVA as specifically indicated in the figure legends. All data are presented as mean ± standard error of the mean (SEM) and values of ∗p<0.05 were considered statistically significant.

## Acknowledgments

Diagrams were created using BioRender. We thank Yuefeng Huang from the Columbia University Medical Center for providing us with a *Nippostrongylus brasiliensis* culture and the Single Cell & Functional Genomics Core for RNA sequencing services.

## Author contributions

Conceptualization: Aleksander Geske, Nathachit Limjunyawong, Xintong Dong, Xinzhong Dong

Methodology: Aleksander Geske, Nathachit Limjunyawong, Xintong Dong

Investigation: Aleksander Geske, Rachel Pan. Celeste Flores

Visualization: Aleksander Geske

Supervision: Nathachit Limjunyawong, Xintong Dong, Xinzhong Dong

Writing—original draft: Aleksander Geske

Writing—review & editing: Aleksander Geske, Rachel Pan, Taylor Follansbee, Xintong Dong, Xinzhong Dong

## Data availability

All data are available in the main text or the supplementary materials. The RNA-sequencing data generated in this study will be deposited in the NCBI Gene Expression Omnibus (GEO) database prior to publication. Processed sequencing results are available in Data S2.

## References

1. Karasuyama, H., Miyake, K., Yoshikawa, S. & Yamanishi, Y. Multifaceted roles of basophils in health and disease. Journal of Allergy and Clinical Immunology 142, 370–380 (2018).

2. Wang, F. et al. A basophil-neuronal axis promotes itch. Cell 184, 422–440.e17 (2021).

3. Voehringer, D. Protective and pathological roles of mast cells and basophils. Nat. Rev. Immunol. 13, 362–375 (2013).

4. Hotez, P. J. et al. Helminth infections: the great neglected tropical diseases. Journal of Clinical Investigation 118, 1311–1321 (2008).

5. World Health Organization. Soil-transmitted helminth infections. https://www.who.int/news-room/fact-sheets/detail/soil-transmitted-helminth-infections (2023).

6. Pullan, R. L., Smith, J. L., Jasrasaria, R. & Brooker, S. J. Global numbers of infection and disease burden of soil transmitted helminth infections in 2010. Parasit. Vectors 7, 37 (2014).

7. Allen, J. E. & Maizels, R. M. Diversity and dialogue in immunity to helminths. Nat. Rev. Immunol. 11, 375–388 (2011).

8. Motomura, Y. et al. Basophil-Derived Interleukin-4 Controls the Function of Natural Helper Cells, a Member of ILC2s, in Lung Inflammation. Immunity 40, 758–771 (2014).

9. Mitre, E. & Nutman, T. B. Basophils, Basophilia and Helminth Infections. in Parasites and Allergy 141–156 (KARGER, Basel, 2005). doi:10.1159/000088886.

10. Sullivan, B. M. & Locksley, R. M. Basophils: A Nonredundant Contributor to Host Immunity. Immunity 30, 12–20 (2009).

11. Sharba, S. et al. Interleukin 4 induces rapid mucin transport, increases mucus thickness and quality and decreases colitis and Citrobacter rodentium in contact with epithelial cells. Virulence 10, 97–117 (2019).

12. Ohnmacht, C. & Voehringer, D. Basophil effector function and homeostasis during helminth infection. Blood 113, 2816–2825 (2009).

13. Galli, S. J. & Tsai, M. Mast cells in allergy and infection: Versatile effector and regulatory cells in innate and adaptive immunity. Eur. J. Immunol. 40, 1843–1851 (2010).

14. McNeil, B. D. et al. Identification of a mast-cell-specific receptor crucial for pseudo-allergic drug reactions. Nature 519, 237–241 (2015).

15. Pundir, P. et al. A Connective Tissue Mast-Cell-Specific Receptor Detects Bacterial Quorum-Sensing Molecules and Mediates Antibacterial Immunity. Cell Host Microbe 26, 114–122.e8 (2019).

16. Dong, X. et al. Neutrophil-Specific Defensin Receptors Prevent Skin Dysbiosis and Bacterial Infection. SSRN Electronic Journal https://doi.org/10.2139/ssrn.3885989 (2021) doi:10.2139/ssrn.3885989.

17. Liu, Q. et al. Sensory Neuron-Specific GPCR Mrgprs Are Itch Receptors Mediating Chloroquine-Induced Pruritus. Cell 139, 1353–1365 (2009).

18. Han, L. et al. A subpopulation of nociceptors specifically linked to itch. Nat. Neurosci. 16, 174–182 (2013).

19. Crowley, G. et al. Quantitative lung morphology: semi-automated measurement of mean linear intercept. BMC Pulmonary Medicine 2019 19:1 19, 206-(2019).

20. Kreider, T., Anthony, R. M., Urban, J. F. & Gause, W. C. Alternatively activated macrophages in helminth infections. Curr. Opin. Immunol. 19, 448 (2007).

21. Wynn, T. Cellular and molecular mechanisms of fibrosis. J. Pathol. 214, 199–210 (2008).

22. Wynn, T. A. & Ramalingam, T. R. Mechanisms of fibrosis: therapeutic translation for fibrotic disease. Nat. Med. 18, 1028 (2012).

23. Schiechl, G. et al. Basophils Trigger Fibroblast Activation in Cardiac Allograft Fibrosis Development. American Journal of Transplantation 16, 2574–2588 (2016).

24. Tsutsui, H. et al. The Basophil-specific Protease mMCP-8 Provokes an Inflammatory Response in the Skin with Microvascular Hyperpermeability and Leukocyte Infiltration. Journal of Biological Chemistry 292, 1061–1067 (2017).

25. Torrero, M. N., Larson, D., Hübner, M. P. & Mitre, E. CD200R surface expression as a marker of murine basophil activation. Clinical & Experimental Allergy 39, 361–369 (2009).

26. World Health Organization. Neglected tropical diseases. https://www.who.int/health-topics/neglected-tropical-diseases?utm_source=chatgpt.com#tab=tab_1 (2024).

27. Craig, J. M. & Scott, A. L. Helminths in the lungs. Parasite Immunol. 36, 463–474 (2014).

28. Babayan, S. A., Read, A. F., Lawrence, R. A., Bain, O. & Allen, J. E. Filarial Parasites Develop Faster and Reproduce Earlier in Response to Host Immune Effectors That Determine Filarial Life Expectancy. PLoS Biol. 8, e1000525 (2010).

29. Huang, L. et al. Eosinophils and IL-4 Support Nematode Growth Coincident with an Innate Response to Tissue Injury. PLoS Pathog. 11, e1005347 (2015).

30. Batugedara, H. M. et al. Hematopoietic cell-derived RELMα regulates hookworm immunity through effects on macrophages. J. Leukoc. Biol. 104, 855–869 (2018).

31. Zunke, F. & Rose-John, S. The shedding protease ADAM17: Physiology and pathophysiology. Biochimica et Biophysica Acta (BBA) - Molecular Cell Research 1864, 2059–2070 (2017).

32. Mizuno, K. et al. Rab27b Regulates Mast Cell Granule Dynamics and Secretion. Traffic 8, 883–892 (2007).

33. Sadhu, C. et al. CD11c/CD18: novel ligands and a role in delayed-type hypersensitivity. J. Leukoc. Biol. 81, 1395–1403 (2007).

34. Zhang, W. et al. The emerging roles of IFIT3 in antiviral innate immunity and cellular biology. J. Med. Virol. 95, (2023).

35. Papavasileiou, S. et al. SinglelJCell Omics Analysis of Human Basophils Reveals Two Transcriptionally Distinct Populations. Allergy https://doi.org/10.1111/all.70209 (2026) doi:10.1111/all.70209.

36. Hamey, F. K. et al. SinglelJcell molecular profiling provides a highlJresolution map of basophil and mast cell development. Allergy 76, 1731–1742 (2021).

37. Miyake, K. et al. Single cell transcriptomics clarifies the basophil differentiation trajectory and identifies pre-basophils upstream of mature basophils. Nat. Commun. 14, 2694 (2023).

38. Wedi, B., Gehring, M. & Kapp, A. The pseudoallergen receptor MRGPRX2 on peripheral blood basophils and eosinophils: Expression and function. Allergy 75, 2229–2242 (2020).

39. Sabato, V. et al. The Mas-Related G Protein-Coupled Receptor MRGPRX2 Is Expressed on Human Basophils and up-Regulated upon Activation. Journal of Allergy and Clinical Immunology 139, AB168 (2017).

40. Toscano, A. et al. Mas-related G protein-coupled receptor MRGPRX2 in human basophils: Expression and functional studies. Front. Immunol. 13, (2023).

