## Supplemental Figures for "A basophil-specific GPCR mediates the immune response to helminth infection"

Aleksander Geske *et al.*

**This PDF file includes:**

Figs. S1 to S4

Fig. S1.

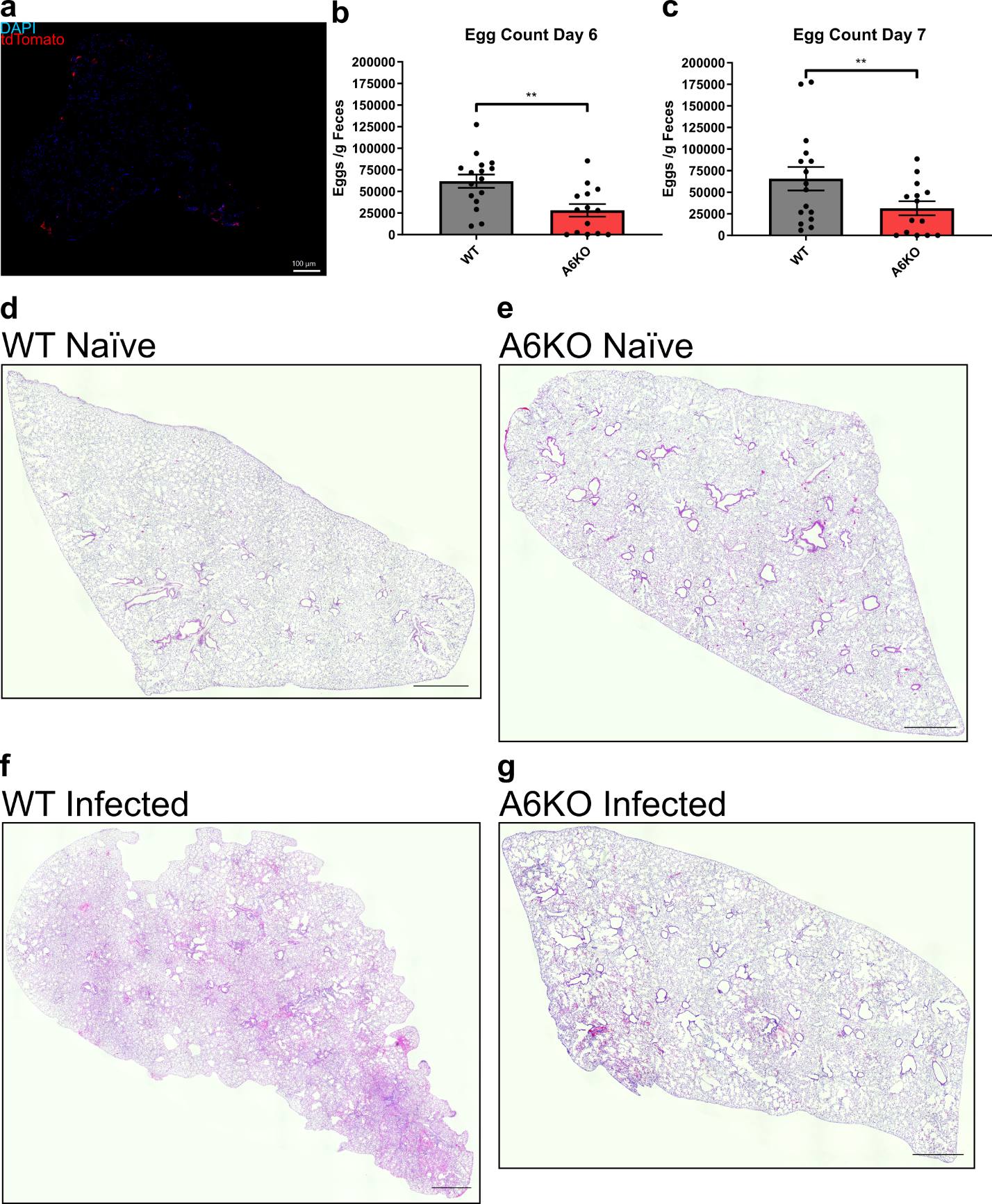

Expanded phenotypic and histological characterization.

(a) Representative confocal images of sections of the L4 DRG from Mrgpra6CreER; Rosa26LSL-tdTomato mice showing DAPI (blue) and tdTomato (red).

(b-c) Day 6 and 7 post-infection egg counts (eggs per gram of feces) from WT and A6KO mice. n = 14-16

(d-g) Representative H&E-stained lung sections from WT and A6KO mice under naïve and day 3 post-infection conditions. Scale bar, 1000 μm.

Results in (b-c) are presented as mean ± SEM from at least three independent experiments. **p < 0.01 (two-tailed unpaired Student’s t test).

Fig. S2.

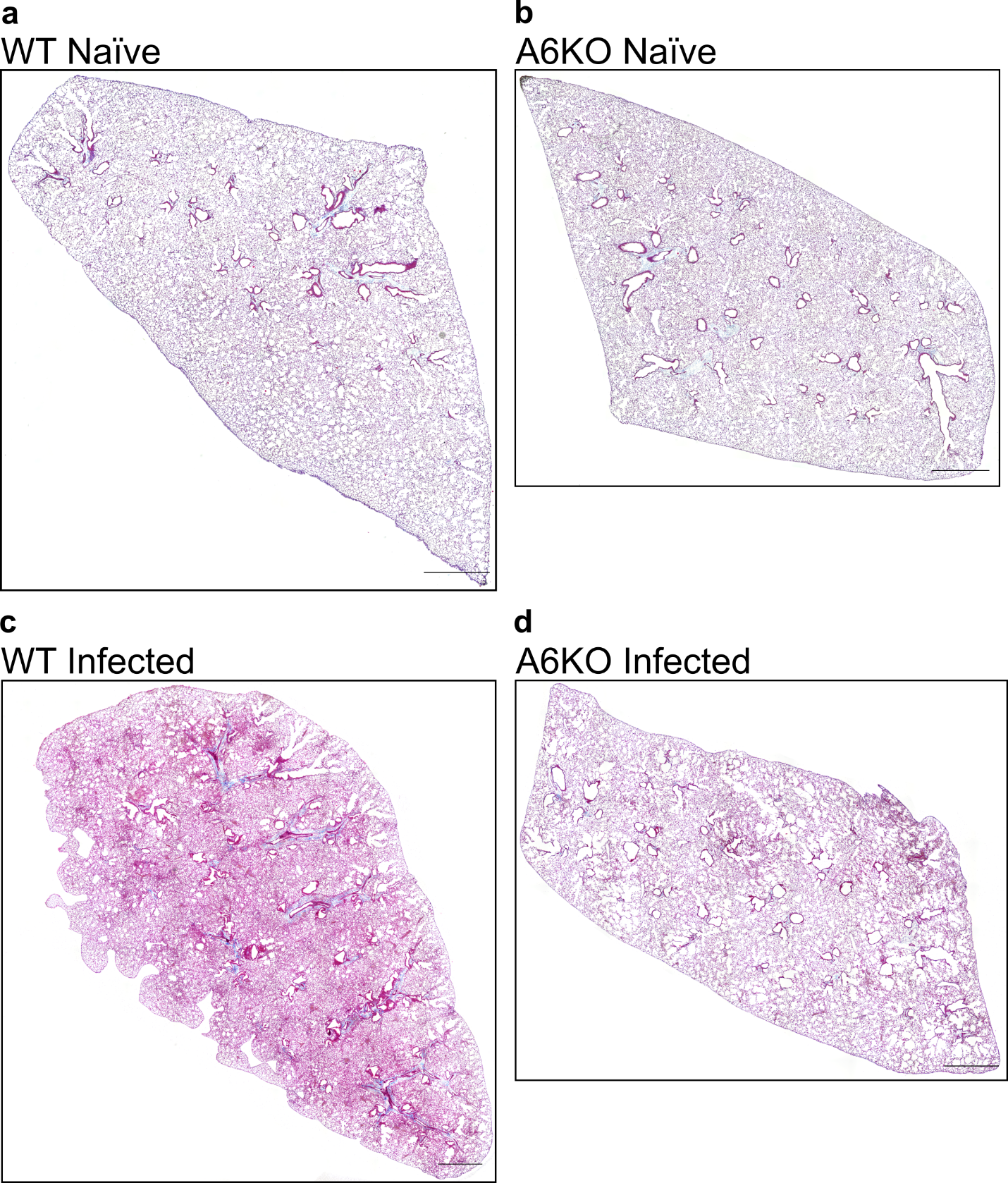

Representative Masson’s trichrome–stained lung sections during early helminth infection.

(a-d) Masson’s trichrome–stained lung sections from WT and A6KO mice under naïve and day 2 post-infection conditions. Images were acquired and processed identically for brightness and contrast. Scale bar, 1000 μm.

Fig. S3.

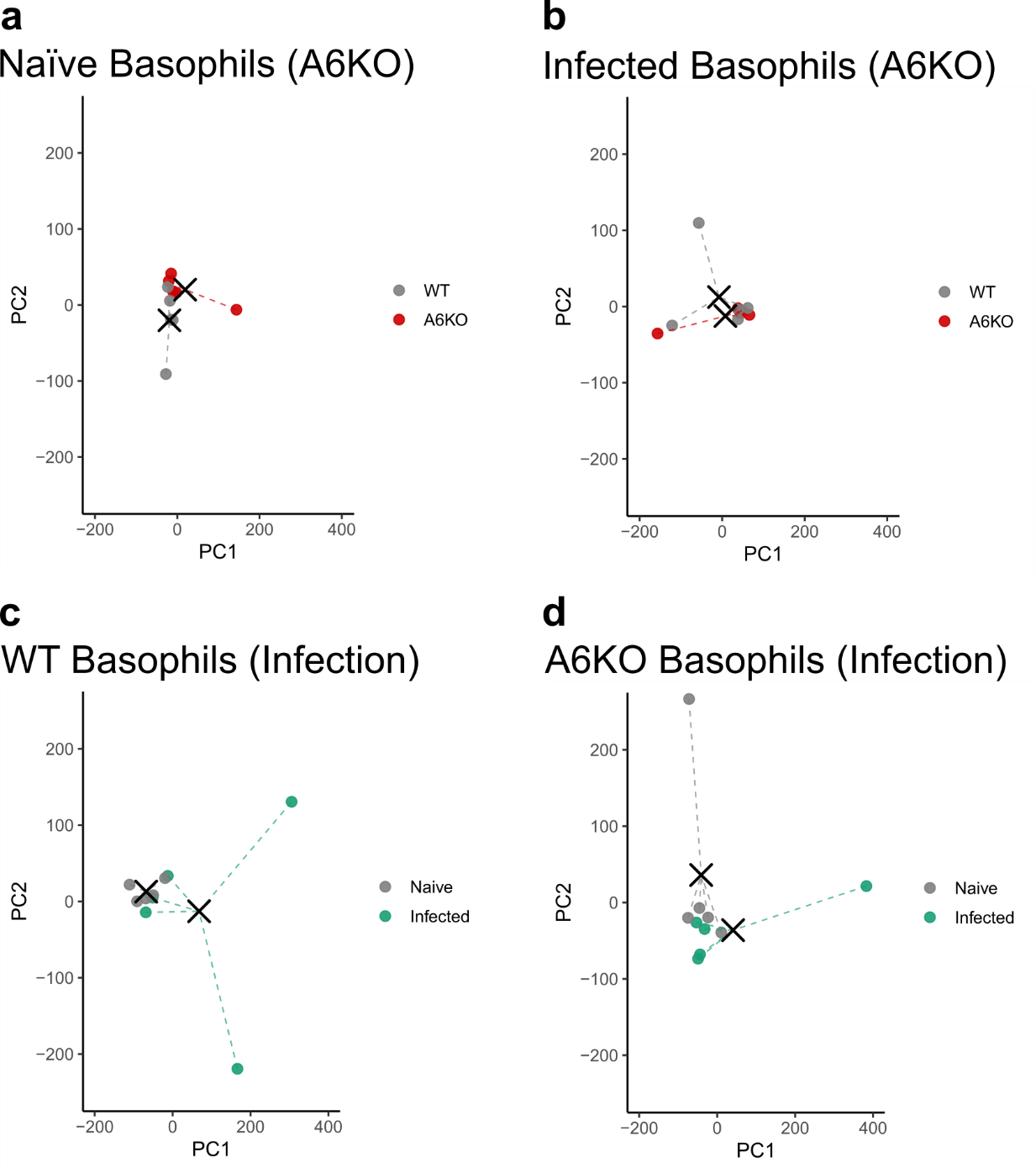

Principal component analysis of basophils during helminth infection.

(a–d) Principal component analysis (PCA) of lung basophils from WT and A6KO mice under naïve and day 3 post-infection conditions. Plots illustrate transcriptional differences associated with A6KO (a, b) or helminth infection (c, d). Black crosses indicate group centroids. Permutation testing of centroid separation revealed significant separation between naïve WT and A6KO basophils (a, p = 0.0356) and a trend toward separation between naïve and infected WT basophils (c, p = 0.0751). No significant separation was observed between infected WT and A6KO basophils (b, p = 0.842) or between naïve and infected A6KO basophils (d, p = 0.423). n = 5 mice per group.

Fig S4.

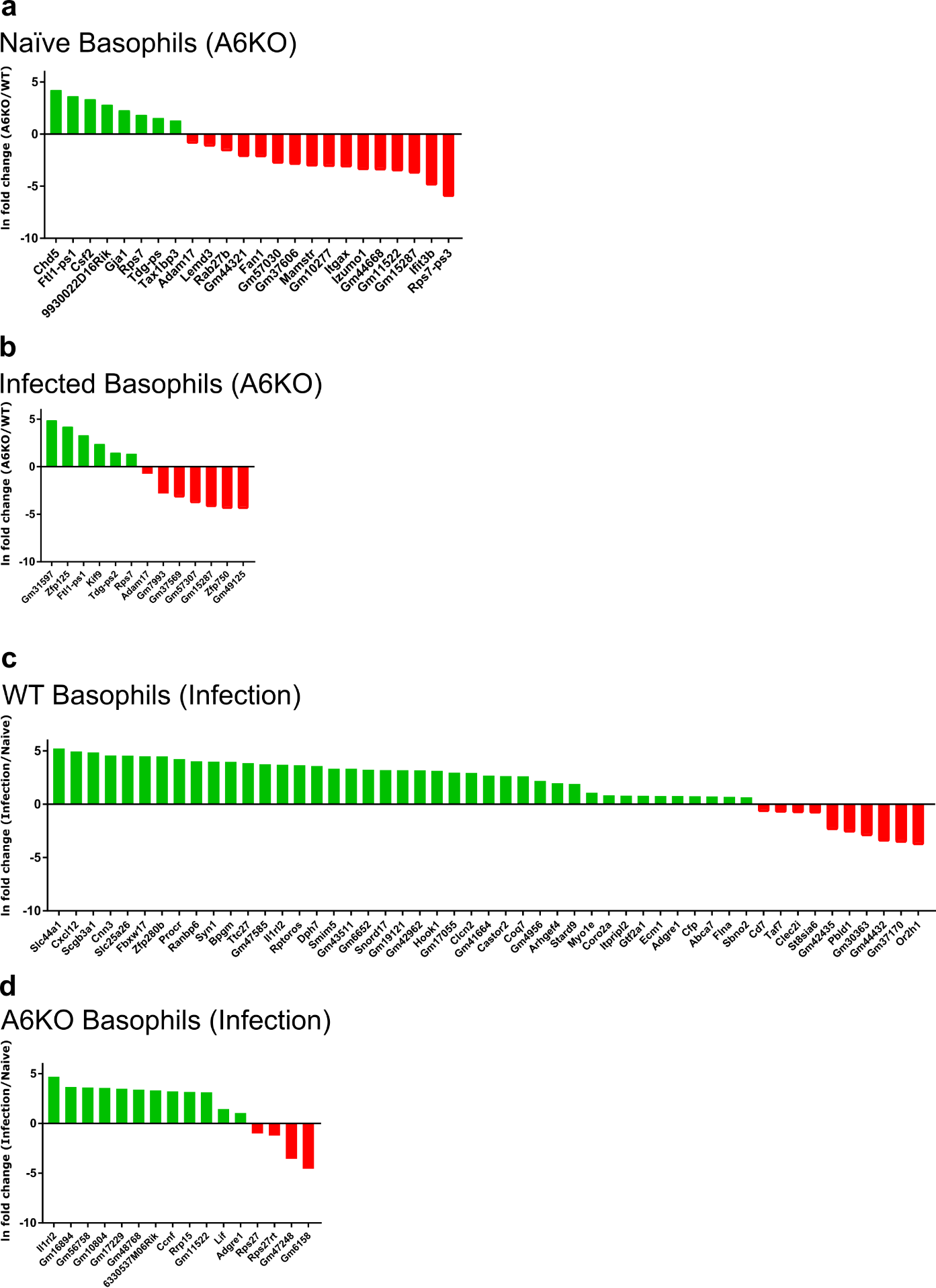

Complete list of significantly differentially expressed genes in basophils

(a-d) Bar graphs showing ln fold change for DEGs in basophils due to Mrgpra6 knockout (A6KO) or N. brasiliensis infection (Infection). Number of DEGs are 24 (a), 13 (b), 51 (c), and 16(d).

Differentially expressed genes were defined using an FDR-adjusted q-value < 0.10, n = 5 individual mice per genotype per condition (WT Naïve, A6KO Naïve, WT Infected, A6KO Infected)
